# Insights into the secondary and tertiary structure of the Bovine Viral Diarrhea Virus Internal Ribosome Entry Site

**DOI:** 10.1101/2021.05.13.444024

**Authors:** Devadatta Gosavi, Iwona Wower, Irene K. Beckmann, Ivo L. Hofacker, Jacek Wower, Michael T. Wolfinger, Joanna Sztuba-Solinska

**Author notes:** To whom correspondence should be addressed J.S.S., Correspondence may also be addressed to M.T.W.

## Abstract

The Internal <u>R</u>ibosome <u>E</u>ntry <u>S</u>ite (IRES) RNA of <u>B</u>ovine <u>v</u>iral <u>d</u>iarrhea <u>v</u>irus (BVDV), an economically significant Pestivirus, is required for the cap-independent translation of viral genomic RNA. Thus, it is essential for viral replication and pathogenesis. We applied a combination of high-throughput biochemical RNA structure probing (SHAPE-MaP) and *in silico* modeling approaches to gain insight into the secondary and tertiary structures of BVDV IRES RNA. Our study demonstrated that BVDV IRES RNA forms in solution a modular architecture composed of three distinct structural domains (I-III). Two regions within domain III are engaged in tertiary interactions to form an H-type pseudoknot. Computational modeling of the pseudoknot motif provided a fine-grained picture of the tertiary structure and local arrangement of helices in the BVDV IRES. Furthermore, comparative genomics and consensus structure predictions revealed that the pseudoknot is evolutionarily conserved among many Pestivirus species. These studies provide detailed insight into the structural arrangement of BVDV IRES RNA H-type pseudoknot and encompassing motifs that likely contribute to the optimal functionality of viral cap-independent translation element.

## INTRODUCTION

<u>B</u>ovine <u>v</u>iral <u>d</u>iarrhea <u>v</u>irus (BVDV) is a member of the genus Pestivirus, family Flaviviridae, that includes the causative agents of economically significant diseases of cattle, pigs, and sheep. The BVDV infections in cattle lead to decreased fertility and milk production, slow fetal growth, diarrhea, respiratory symptoms, reproductive and immunological dysfunction (Khodakaram-Tafti and Farjanikish 2017). According to the ICTV virus taxonomy profile (Simmonds et al. 2017), two main genotypes of BVDV, namely type 1 of low-virulence and type 2 of high-virulence, have been recognized and are estimated to cause $20 and $57 million in total annual losses, respectively (Houe 1999). Besides the two BVDV genotypes, several genetic isolates have been distinguished, within BVDV type 1 (Vilček et al. 2001; Burks et al. 2011a).

The BVDV genome is a positive-sense single-stranded RNA (12.5 kb) that codes for one open reading frame (ORF). The coding region is preceded by the distinctly structured 5’ untranslated region (5’ UTR), which folds into the Internal <u>R</u>ibosomal <u>E</u>ntry <u>S</u>ite (IRES) (Eterhans et al. 2010). By definition, IRES is the RNA domain that recruits ribosomes to the internal region of mRNAs to initiate translation through a cap-independent pathway. IRES domains can be found in the genomic RNAs of many pathogenic viruses. In Hepatitis C virus (HCV) genomic RNA, it mediates the initiation of translation by recruiting the subset of canonical translation initiation factors (eIF2, eIF3), methionine tRNA (Met-tRNAi), and by orienting the 40S ribosomal subunit at the translational initiation codon(Tsukiyama-Kohara et al. 1992; Rijnbrand et al. 1995). Cricket Paralysis Virus (CrPV) IRES RNA directs ribosomes without the requirement of additional translational factors (Wilson et al. 2000), while IRES RNAs of Poliovirus (PV) (Kühn et al. 1990) and Foot and Mouth disease virus (FMDV) (Pelletier and Sonenberg 1988) rely on binding of eIFs, Met-tRNAi, and additional proteins referred to as the IRES trans-acting factors (ITAFs) (Burks et al. 2011a; Thakor and Holcik 2011; Martinez-salas 2018; Burks et al. 2011b)

According to previous structural studies, the viral IRES RNAs fold into intricate secondary and tertiary arrangements that provide a structural scaffold for cellular initiation factors and trigger conformational changes in the 40S ribosomal that drive translation initiation by active mechanism (Komar and Hatzoglou 2015; Ghassemi et al. 2017; Yu et al. 2011; Yamamoto et al. 2015). For example, the HCV IRES consists of four structurally defined domains (I-IV), with domain II inducing the open conformation of the 40S subunit and III and IV forming a functionally essential pseudoknot (Lukavsky 2009). Pseudoknots are formed upon base-pairing of a single-stranded region of RNA in the loop of a hairpin or a bulge to a stretch of complementary nucleotides elsewhere in the RNA chain. This folding strategy can generate a vast number of three-dimensional folds, which exhibit a diverse range of highly specific functions (Brierley et al. 2007). CrPV IRES also displays a multi-domain composition, with the residues of one domain supporting a pseudoknot. In the case of BVDV IRES (SD-1 isolate, genotype 1), thermodynamic and phylogenetic studies suggested that its translational functionality is also controlled by the formation of a pseudoknot (Brierley et al. 2007; Moes and Wirth 2007). Burks et al. (2011), using comparative structural studies, predicted that the BVDV IRES pseudoknot involves interaction between the loops of hairpin 3 and 4 within domain III (Burks et al. 2011a; Moes and Wirth 2007; Pestova and Hellen 1999). Toeprinting *in vitro* analysis involving BVDV IRES, 40S rRNA, and eIFs suggested that the ribosome makes multiple contacts with the BVDV IRES. Similar observations have been made for HCV IRES RNA (Fukushi et al. 2001; Otto et al. 2002; Laletina et al. 2006; Babaylova et al. 2009), outlining particularly interesting contacts existing between 18S rRNA and the pseudoknot formed within IRES domain III (Yamamoto et al. 2015; Pestova and Hellen 1999). Furthermore, the presence of an RNA pseudoknot in BVDV IRES RNA was supported by compensatory mutations, which restored translation (Moes and Wirth 2007).

In this study, we investigated the secondary structure of BVDV IRES RNA using selective 2′-hydroxyl acylation analyzed by primer extension and mutational profiling (SHAPE-MaP) (Smola et al. 2015a, 2015b). The analysis, which has been performed on two *in vitro* synthesized constructs which contain all essential elements supporting the cap-independent translation, revealed that BVDV IRES RNA folds into three distinct structural domains (I-III). Two single-stranded regions of domain III were shown to support the formation of an H-type pseudoknot, thus corroborating earlier studies (Le et al. 1995). To obtain a better picture of pseudoknot formation, we performed RNA 3D simulations with ernwin (Kerpedjiev et al. 2015) and SimRNA (Boniecki et al. 2015) on a trimmed construct of BVDV1-NADL IRES. Tertiary structure simulations confirmed that an H-type pseudoknot can be formed under the condition that two adjacent helices are oriented in a well-defined range of angles against each other. On a broader scale, we performed a comparative genomics screen of the 5’ UTR in the genus Pestivirus, which confirmed pervasive evolutionary conservation of the IRES region, and provided evidence for the presence of a pseudoknot in all studied viruses.

## RESULTS

### The secondary structure of BVDV IRES RNA

SHAPE-MaP is a high-throughput biochemical probing technique that can be used to characterize RNA secondary structure at a single-nucleotide resolution (Supplementary Figure S1). In SHAPE-MaP, unpaired nucleotides (nts) are acylated more readily at the 2′-hydroxyl ribose moieties by an electrophilic reagent, e.g., 1-methyl-7-nitroisatoic anhydride (1M7) (Mortimer and Weeks 2007), than those that are paired, and the reactivity biases are employed to address RNA secondary structure (Sztuba-Solinska et al. 2017). The modified RNA is directed to reverse transcription reaction in the presence of Mn^2+^ ions that induce reverse transcriptase to read through the bulky 2’-O-adducts and insert mutations (Siegfried et al. 2020). The resulting cDNA products carrying the mutations are directed to stepwise amplification to generate DNA libraries directed to next-generation sequencing.

We performed structural probing of BVDV IRES RNA on two *in vitro* synthesized transcripts; the short transcript (560 nts) included the BVDV IRES RNA sequence only; the long transcript (1296 nts) additionally contained the EGFP cassette (nts 561 - 1296), and the MS2 hairpin (nts 523 - 543) at its 3’ end. The IRES RNA constructs used in this study were derived from BVDV-NADL genomic RNA (Supplementary Figure S2), and contain RNA sequences, that were outlined as necessary for supporting the expression of EGFP protein by the cap-independent translation (Ghassemi et al. 2017; Ghaderi et al. 2014). We confirmed the homogeneous conformation of both folded RNAs by native gel electrophoresis prior to structural probing (Supplementary Figure S3). The obtained reactivity values were used to classify probed residues into four groups represented by low (< 0.25), medium (0.25 - 0.80), high reactivity (0.8 - 2), and hyperreactivity (> 2). The comparison of SHAPE reactivity profiles for short and long IRES RNA constructs performed by custom R-scripts, showed a positive correlation established by an average Pearson correlation coefficient of 0.8398 (*p*-value = 0.022) (Supplementary Figure S4).

Our analysis revealed that the BVDV IRES structure can be divided into three domains labeled I-III, with individual stem-loops (SL) within each domain being assigned letter codes (Figure 1). Domain I contains three SLs; SL Ia is enclosed within nts 8-39 and includes a 19 nt apical loop with an equal number of purines and pyrimidines. SL Ib involves nts 41-77 and has a purine-rich apical loop consisting of 14 nts and a stem with unpaired cytosine at position 42. SL Ic has an adenine-rich 19 nts apical loop, and a stem that is closed with a non-canonical GU wobble base pair and another wobble base pair at nt position 73 and 107 within the stem.

**Figure 1.**
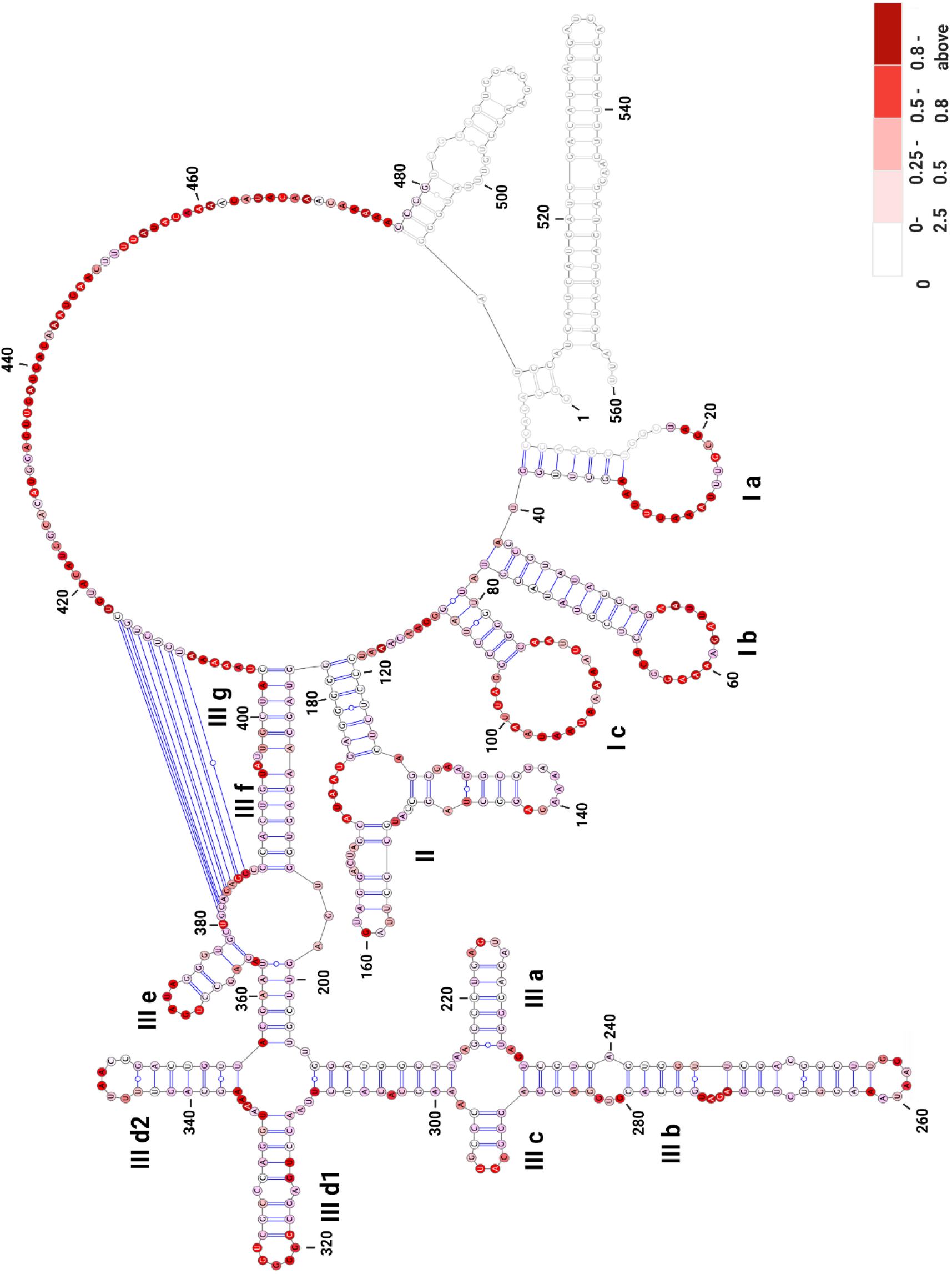
The secondary structure model of BVDV IRES RNA derived from SHAPE-MaP probing. The three main domains are labeled (I-III) with outlined individual RNA motifs: SL Ia, SL Ib, SL Ic, SL II, SL IIIa, SL IIIb, SL IIIc, SL IIId1, SL IIId2, SL IIIe, SL IIIf, SL IIIg. The single-stranded residues at positions 381-387 and 410-416 of domain III are involved in the formation of an H-type pseudoknot. The residues lacking reactivity values correspond to the sites of primer annealing (light grey). The secondary structure is color-coded according to the SHAPE reactivity values.

Domain II includes nts 119-182 and consists of two SLs interconnected by a three-way junction (nts 125-132 and 170-176), and the main stem that includes a GU wobble base pair (nts 122 and 179). The 5’ SL includes nts 127-149 that fold into two short stems interrupted by an asymmetrical internal loop with three nts at the 5’ side and mispaired adenine at the 3’ side. The apical loop includes predominantly adenines, which are largely not reactive to SHAPE reagent, likely due to extensive base stacking interactions characteristic for A-rich sequences (Hurst et al. 2018). The 3’ SL extends from nts 153-170 and contains a 3’ four nts bulge and a UNRN tetraloop (UAGU), where N stands for any nucleotide and R is a purine base. These types of tetra-loops are frequently present in apical loops involved in the U-turn formation, where the backbone changes direction immediately following the initiating U, and the Watson–Crick faces of the bases immediately 3′ of the U are exposed to the solvent (Stallings and Moore 1997; Moore 1999).

Domain III (nts 183-403) includes numerous internal stems and loops; the main stem is formed by SL IIIf and IIIg, interrupted by a 2 nt bulge, followed by a three-way junction that connects SL IIIe and the apical IRES portion. Successively, a four-way junction connecting SL IIId1 and IIId2 is present, followed by a four-way junction connecting SL IIIa, b, and c. The stem of SL IIIa includes a GU wobble pair and an AGUA tetra-loop with less-reactive UA residues (Stallings and Moore 1997). The SL IIIb forms the longest hairpin of this region, closed with a purine-rich apical loop (nts 256-261). It includes an asymmetrical internal loop at its 3’ side, a downstream 3 nt bulge, and a cytosine mismatch at positions 250 and 267. Two GU wobble base pairs are involved in the formation of the IIIb stem. The SL IIIc includes three canonical base pairs and a CAUG tetra-loop belonging to YNMG tetra-loop family, where Y stands for U/C, N, and M is for A/C/U. Previous NMR, circular dichroism, and functional group substitution studies in 16S rRNA indicated the unusual thermodynamic stability of YNMG tetra-loop and showed its involvement in tertiary interactions (Proctor et al. 2002). The SL IIId1 and IIId2 contain eight and seven base pairs, respectively. SL IIId1 is closed with an apical penta-loop that consists of highly reactive G residues (median SHAPE reactivity of 1.020). This type of G-rich apical loop was found in the 5’ UTR SL2 and SL3 of the HIV-1 genome involved in binding the nucleocapsid protein (Paoletti et al. 2002). The SL IIId2 involves a stem with two GU wobble base pairs and a 6 nt apical pyrimidine-rich loop. The SL IIIe includes nts 364-378 and is closed with UGAUA penta-loop. The most distinctive structural feature of domain III is the involvement of single-stranded residues at positions 381-387 (5’-GCAGAGG-3’, median SHAPE reactivity of 0.617) and 410-416 (5’-UCUCUGC-3, median SHAPE reactivity of 0.115) in the formation of an H-type pseudoknot. The single-stranded region immediately downstream of the pseudoknot harbors the AUG codon (nts 429-431). We noted that numerous residues within BVDV IRES RNA display hyperreactivity towards SHAPE reagents (Supplementary Table S4, Supplementary Figure S4). These are found within the loops (nts 26, *22*, 54, 58, 60, 9*1*, 99), bulges (nts 271, 273), single-stranded regions (nts 116, 438, 444, 455, 461,465, 469), and near the pseudoknot (nts 409, 417) (Supplementary Figure S5).

We used SuperFold to compute a SHAPE-MaP assisted minimum free energy structure as well as base pair probabilities of the BVDV IRES RNA by partition function folding. The obtained model was visualized as a color-coded arch plot (Figure 2). Well-defined pairings outlined by high pairing probability between 80 - 100% (green arcs) are visible in regions folding into SL Ia - c and SL IIIa - e. Regions with low pairing probabilities < 30% (yellow and grey arcs) are scarce and dispersed among residues, that per SHAPE modeling, are predicted to be single-stranded. Overall, the arc model displayed strong agreement between nucleotide-resolution experimental SHAPE reactivity data and modeled base-pairing. We also assessed how accurately the secondary structure motifs are defined by their sequence and the probing data by calculating the Shannon entropies at nucleotide resolution (Siegfried et al. 2020; Huynen et al. 1997). We noted that the region corresponding to the Domain III (nts 200 – 380) displays low SHAPE reactivity values, low Shannon entropy values, and high pairing probability, which reflect likely reflects its structural and functional significance. On the other hand, the regions that display low (nts 130 - 200) and high (nts 420 – 490) SHAPE reactivity values and high Shannon entropy correspond to Domain II and single-stranded residues of Domain III that are predicted to form a pseudoknot (nts 381-387 and 410-416). These parameters were shown previously to be characteristic of multiple and/or interconverting RNA structures (Huber et al. 2019; Mauger et al. 2015).

**Figure 2.**
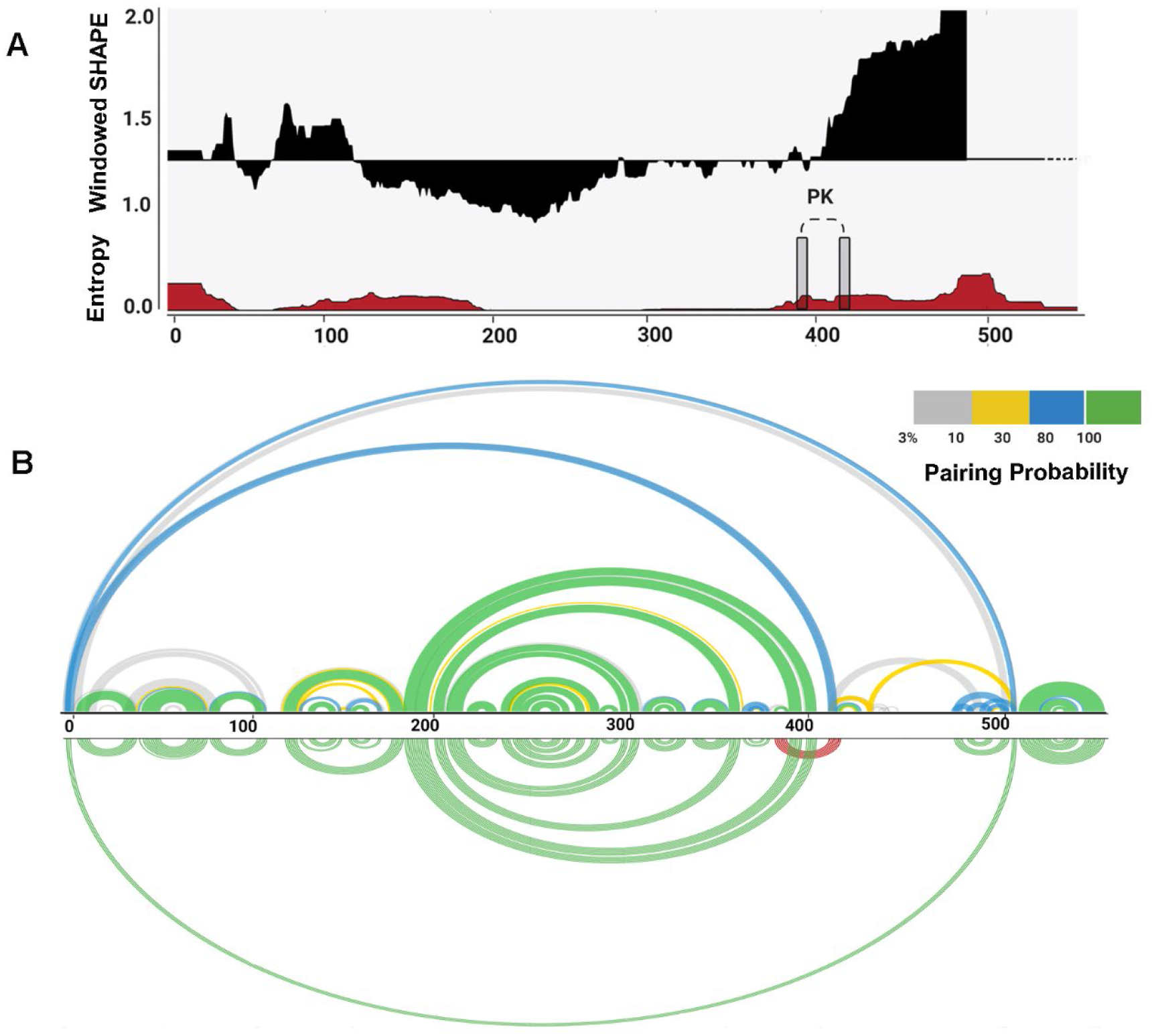
Structural features of BVDV IRES RNA. (**A**) Windowed SHAPE reactivities (black) and Shannon entropies (brown) are indicated. The X-axis represents the nucleotide position, and Y-axis corresponds to windowed SHAPE reactivities (top) and entropy values (bottom). Single-stranded regions involved in pseudoknot formation are indicated with grey rectangles. (**B**) Arc plots representing the base-pairing probabilities for BVDV IRES RNA. The top arc plot is color-coded according to pairing probability scale. The bottom arc plot includes the predicted H-type pseudoknot (red).

### Evolutionary conservation of pseudoknot in the genus Pestivirus

To obtain deeper insight into the evolutionary conservation of the BVDV IRES region, and in particular conservation of the pseudoknot, we performed a comparative genomics screen within the genus Pestivirus. To this end, we searched for hits of the Rfam Pestivirus IRES (RF00209) covariance model among 13 representative Pestivirus genomes resulting in high-quality (high bit score) hits for each input sequence. Multiple sequence alignment (Supplementary Figure S6), and consensus structure prediction of these structurally homologous elements revealed rich covariation patterns throughout the entire IRES region (Figure 3). A test for statistically significantly covarying base pairs with R-scape reported 23 hits, one of them in the pseudoknot.

**Figure 3.**
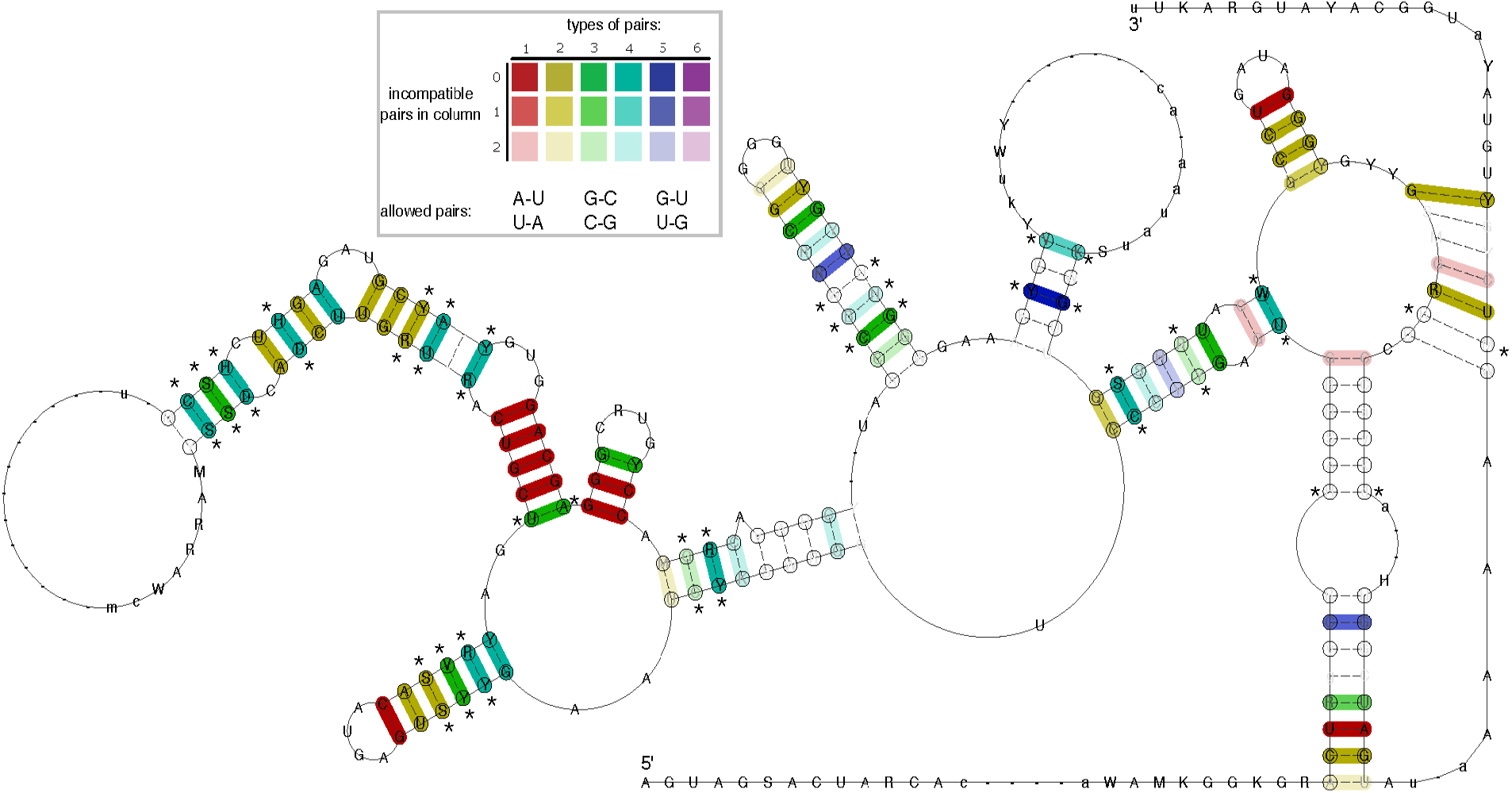
The consensus of IRES Domain III secondary structure prediction generated for 13 Pestivirus genomes highlights rich covariation patterns by the different coloring of base pairs, following the RNAalifold color scheme. Red indicates nucleotide-level sequence conservation, while other colors highlight increasing covariation levels by structure-conserving nucleotide substitutes, as depicted in the insert. The model includes nucleotides with a frequency greater than the average in the underlying alignment in IUPAC notation. Dashes along the consensus sequence indicate positions where the majority of sequences had gaps. Circled nucleotides highlight compensatory mutations, i.e., cases where both nucleotides of a base pair are mutated, such as UA -> GC. Statistical significantly covarying base pairs are marked with an asterisk.

### The modeling of BVDV IRES RNA 3D structure

SHAPE-assisted secondary structure prediction reported the existence of an H-type pseudoknot near the basal stem of Domain III. By analyzing the predicted secondary structure, one might expect a co-linear arrangement of SL IIIf and IIIg, resulting in coaxial stacking of the two helices. However, under such a scenario, spatial separation of the complementary pseudoknot interaction regions induced by the putatively coaxially stacked helical stems SL IIIf and SL IIIg might be too large to form the pseudoknot. As RNA is known to adopt a highly specific yet flexible tertiary architecture to perform its biological functions, we were intrigued by the question of whether or not this pseudoknot interaction is sterically feasible in 3D.

To gain more insight into the tertiary structure of BVDV IRES, we performed coarse-grained RNA 3D simulations with a purposefully trimmed construct that encompasses relevant parts of Domain III that are involved in pseudoknot formation. Specifically, 5’ and 3’ portions of the original construct were truncated after positions 179 and 425, respectively. Likewise, the four-way junction that contains SL IIId1, SL IIId2, and the substructure comprising SL IIIa, b, and c, was retained as an apical loop element with a connected backbone. This construct, denoted BVDVsegment_180_425 (Supplementary Figure S7), features the same structural traits as the original full-length construct. In particular, pKiss single sequence folding predictions suggest the formation of an H-type pseudoknot as expected. Likewise, this truncated construct represents a reasonably small system that allows for detailed tertiary structure analysis *in silico*.

We used ernwin to test whether the proposed secondary structure, encompassing the H-type pseudoknot, can be embedded in 3D. Operating on secondary structure elements, ernwin employs a coarse-grained sampling approach and distinguishes between stems, bulges, and various loop types, each represented by cylinders that fit the helix axis. 3D structures are built from fragments of already known RNAs, and tertiary structures are guaranteed to always match the input secondary structure. Building on this concept, we constructed initial 3D models of BVDVsegment_180_425, thereby confirming that the proposed H-type pseudoknot is sterically feasible.

In parallel, we set out to predict the tertiary structure of BVDVsegment_180_425 with SimRNA. This Monte Carlo sampler uses a knowledge-based potential and achieves coarse-graining by reducing each nucleotide to five beads. We performed massively parallel replica-exchange Monte Carlo simulations to sample the low-energy portion of the underlying energy landscape, using secondary structure restraints that ensure the formation of all helices predicted for BVDVsegment_180_425, including the pseudoknot. A snapshot of a low-energy conformation is shown in (Figure 4). Clustering of predicted low-energy structures allowed us to characterize two sets of highly compact target structures, 38 in total, that differ in the size of the bulge between helical segments IIIf and IIIg, i.e., 2 nt in set 1 (17 candidate structures) and 3 nt in set 2 (21 candidate structures). The sets of candidate structures were then subjected to follow-up analysis with forgi, focusing on the angles between the pseudoknot helix and SL IIIf (angle β) and SL IIIf and SL IIIg (angle α). As shown in (Figure 5), both angles are distributed in well-defined intervals for all 38 low-energy structures. While α falls between 37 and 149 degrees, β is dispersed between 129 and 174 degrees. This suggests that all 3D candidate structures are well-defined and that there is nearly quasi-coaxial stacking between the pseudoknot helix and SL IIIf. Likewise, our data indicate that there is a marked twist between SL IIIf and IIIg. Notably, structures with a 2 nt bulge are less arched than those with a 3 nt bulge, which is in good agreement with their expected behavior. An important finding of this analysis is that the pseudoknot can only be formed when SL IIIf and SL IIIg are oriented at an acute angle, facilitated by the central bulge.

**Figure 4.**
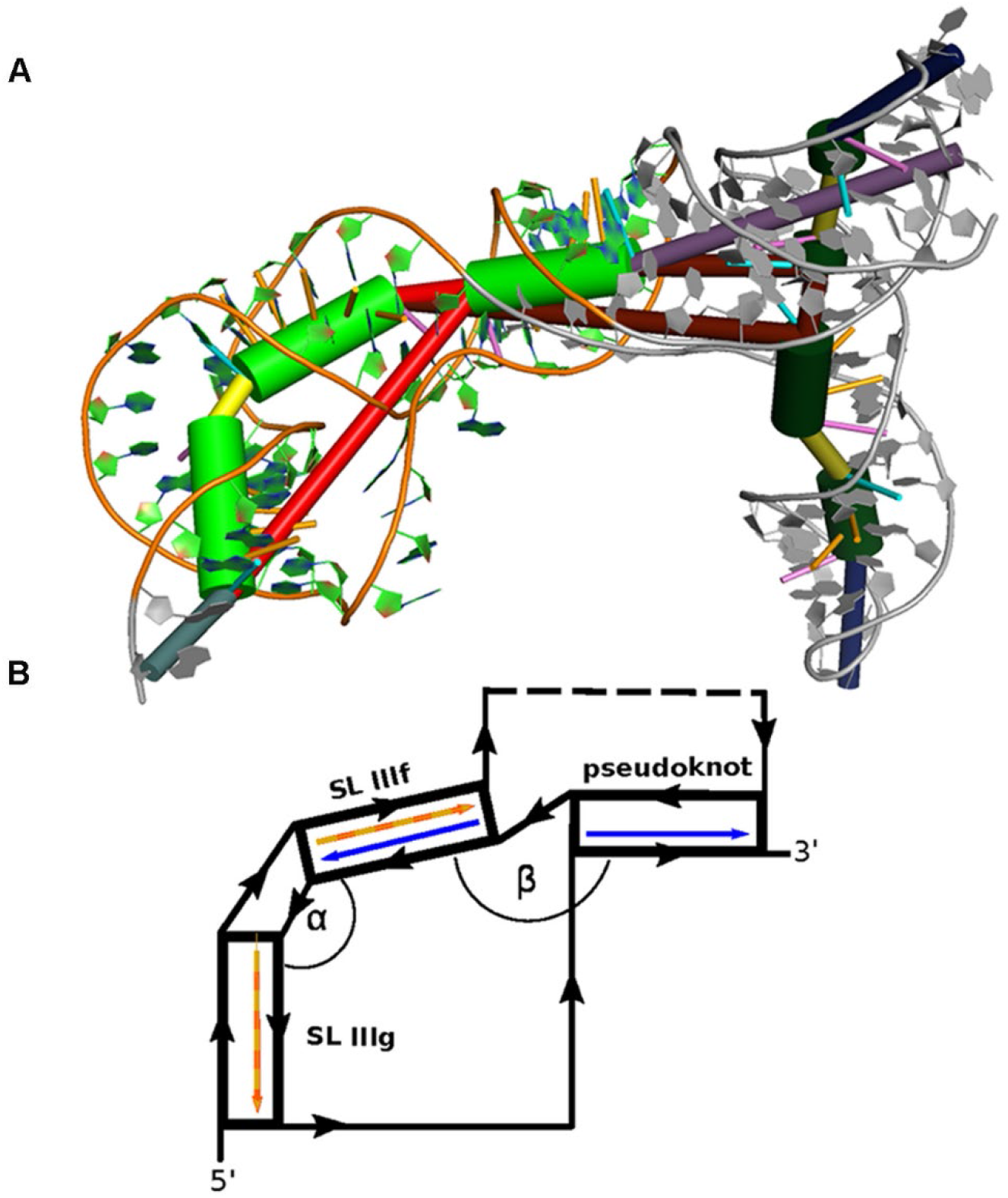
(**A**) Rendering of a low-energy tertiary structure snapshot of the truncated construct BVDVsegment_180_425 overlaid with coarse-grained 3D elements. Green cylinders highlight helical regions, with the rightmost representing H-type pseudoknot, and the middle and leftmost green cylinders portraying stem-loops IIIf and IIIg, respectively. The bulge between SL IIIf and IIIg is depicted in yellow, and the multiloop enclosed by the formation of the pseudoknot is plotted as a long red connection. The short red connection between the pseudoknot helix and SL IIIf represents a single unpaired C between these helices. Other structural elements are greyed out. The quasi-colinear arrangement of the pseudoknot and SL IIIf is noticeable, while the bulge induces a marked twist between SL IIIf and IIIg. (**B**) Schematic representation of the three helices encompassing the pseudoknot, SL IIIF, and IIIf, sketching their approximate spatial arrangement. Angles between adjacent helical segments are defined by stem vectors, i.e., angle α between SL IIIf and SL IIIg (orange and red vectors), and β between SL IIIf and the pseudoknot helix (blue vectors). The directionality of the backbone is indicated by black arrows. Regions of BVDVsegment_180_425, which are not directly involved in the spatial arrangement of these helices, are depicted as a dashed segment along the backbone.

**Figure 5.**
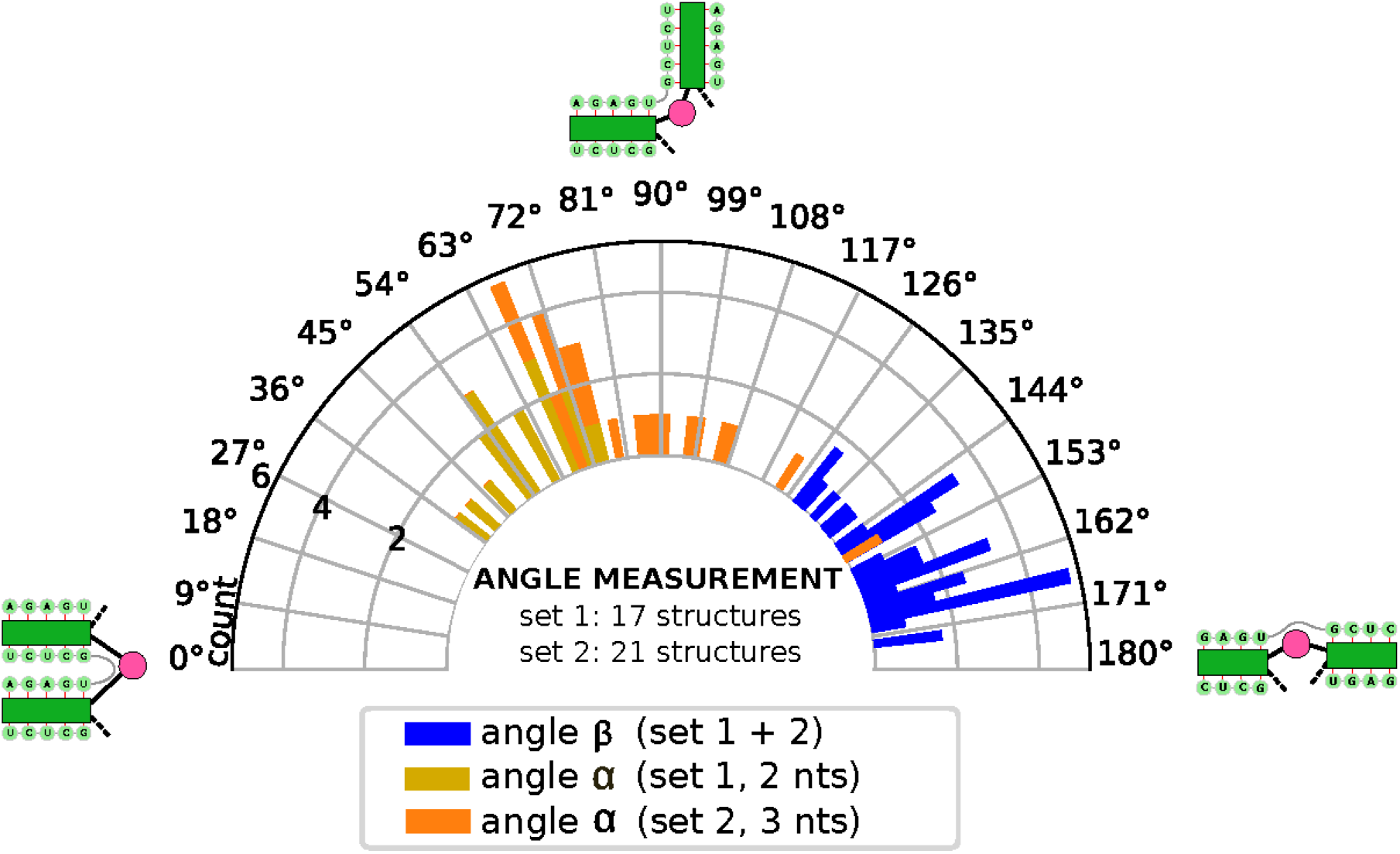
Distribution of angles α and β in the low-energy 3D structure of BVDVsegment_180_425. The relative orientation of helices was assessed by a helix-centered representation of RNA 3D structure. As stretched by the relative geometry of green helical segments, angles close to 0° account for parallel stems, whereas angles close to 180° indicate colinear arrangement and can be indicative of coaxial stacking. Analysis of candidate structures from SimRNA simulations of BVDVsegment_180_425 shows that angles between helices are distributed in well-defined intervals. Set 1 and set 2 comprise low-energy structures predicted by SimRNA, whose helices SL IIIf and IIIg are separated by a bulge of 2 and 3 nts, respectively. For set 1, α lies between 37 and 75 degrees, while for set 2, values are distributed between 66 and 149 degrees. This is expected, as a larger bulge allows for more conformational flexibility of the two adjacent helices. Contrary, β falls in a range between 130 and 174 degrees for set 1 and 129 and 173 degrees for set 2. Values for both sets were combined and depicted in blue.

## DISCUSSION

RNA viruses are known to use exceptionally thrifty molecular strategies to optimize their replication cycle. They often contain functional RNA motifs within non-coding regions that control crucial steps of their infectivity cycle and structural switches that aid their regulation. The IRES RNA within the 5’ UTRs of BVDV is one of such complex structural domains known to govern the efficiency of cap-independent translation (Pestova and Hellen 1999). Currently, there are no high-resolution secondary or tertiary structural data available for BVDV IRES RNA (Burks et al. 2011a). In this study, we applied the SHAPE-MaP secondary structure probing method and computational 3D modeling to gain insight into the secondary and tertiary structure of BVDV IRES RNA.

The biochemical probing analysis of two BVDV IRES constructs resulted in the secondary structure model that shares a significant correlation between RNAs. Overall, BVDV IRES RNA folds into three major structural domains (I – III), each containing well-defined motifs, i.e., hairpins, stems, interspersed among less constrained regions, i.e., multi-way junctions, and loops. The structure includes some hyperreactive residues primarily contained within flexible single-stranded regions. These residues were previously reported as accommodating a C2-endo ribose conformation (McGinnis et al. 2012). This conformation of ribose shows slow dynamics and favors intra-residue H bonding between the 2′-hydroxyl and the 3′-oxygen, enhancing the reactivity of the 2′-hydroxyl towards SHAPE reagent (Vicens et al. 2007). Previously, the 16S rRNA of *Thermus thermophilus* was shown to include a residue that displayed hyperreactivity towards alkylating probing reagent, and that was critical for the structure and functionality of a pseudoknot (Gregory and Dahlberg 2009).

The key finding relates to the most structurally well-defined domain III (low SHAPE reactivities, low Shannon entropy values, high pairing probability), which includes two single-stranded regions that support the formation of an H-type pseudoknot. The 3D modeling of the BVDV IRES RNA provides further insight into the structural arrangements of individual motifs, predicting quasi-coaxial stacking between SL IIIf and the pseudoknot helix. These coaxial stackings can be critical for stabilizing the IRES pseudoknot and can be formed only if the angle between SL IIIg and SL IIIf is acute. Similar stabilization processes have been shown for T2 bacteriophage mRNA of gene 32 (Holland et al. 1999), turnip yellow mosaic virus (TYMV) (Dumas et al. 1987), and simian retrovirus-1 (SRV-1) RNA pseudoknots (Sung and Kang 1998). The large junction connecting the individual RNA motifs of domains I – III likely provides a suitable movement and intrinsic flexibility to the single-stranded regions within domain III, which support the formation of a pseudoknot.

Previously, the formation of a pseudoknot in BVDV IRES RNA has been proposed based on thermodynamic predictions and Monte Carlo simulations (Le et al. 1995). The comparative phylogenetic analysis performed on BVDV IRES RNA sequences from genotypes 1 and 2 provided further evidence supporting the formation of a pseudoknot (Burks et al. 2011a). These studies showed that the pseudoknot engages between four and seven nucleotides, depending on the BVDV strain, with residues 5’-CAGA-3’ and 5’-UCUG-3 being shared by all isolates. Here, we performed a computational screen of the IRES Domain III in more than a dozen phylogenetically related viruses. Consensus structure prediction revealed rich covariation support in this evolutionarily conserved region of the Pestivirus IRES. Importantly, we could identify statistically significant covariation in 22 base pairs, including the pseudoknot, thus providing computational evidence for the crucial role of the pseudoknot in IRES functionality.

Site-directed mutagenesis analysis indicated that BVDV IRES RNA constructs with impaired base-pairing within these residues lost their translational activity, while compensatory mutations reversed that effect (Moes and Wirth 2007). Furthermore, mutational and structural studies suggested the formation of structurally similar pseudoknots in the IRES of CSFV and HCV RNAs (Fletcher and Jackson 2002; Wang et al. 1995). Additionally, it has been shown that the BVDV and HCV 5’ UTRs share a considerable extent of sequence and structure similarity (Sung and Kang 1998; Spahn et al. 2001; Le et al. 1996). The shared features include the formation of multiple SLs, which are globally organized into three domains surrounding the centrally positioned pseudoknot (Grassmann et al. 2005), with HCV IRES folding into additional domain IV, positioned externally (Lemon and Honda 1997). Interestingly, in both HCV and BVDV IRES RNAs, domain III includes hairpins IIIa, IIId1, and IIIe, which apical loops, i.e., tetra-loop, G-rich loop, and penta-loop have highly conserved nucleotide sequence (Lukavsky 2009).

Although the relationship between the IRES pseudoknot and initiation codon in BVDV RNA has not been addressed, previous studies performed on HCV IRES RNA indicated that the formation of a pseudoknot leads to the correct positioning mRNA open reading frame (ORF) in the 40S ribosomal binding cleft (Berry et al. 2010). In HCV IRES, it was shown by SHAPE probing that upon binding of the 40S subunit P-site to the AUG codon, the residues at and beyond the start codon show a substantial increase in the flexibility facilitating translation initiation (Berry et al. 2011). Similar structural probing experiments in the presence and absence of ribosomes can be performed to understand the effects of the translational machinery assembly on BVDV IRES RNA. Additionally, various therapeutic strategies involving the identification of small molecule inhibitors, antisense oligonucleotides, ribozymes, which target and disrupt the HCV IRES RNA and prevent the binding of translational factors, abolishing the viral replication, have also been developed (Pawlotsky et al. 2007). Considering the similarity between HCV and BVDV IRES RNA, analogous strategies can be applied for interfering with BVDV replication.

## MATERIAL AND METHODS

### Cloning IRES from BVDV-NADL into pIW-IRES(eMS2hp)-EGFP expression vector

Oligonucleotide primers used in the study were purchased (IDT) and are listed in (Supplementary Table S1). All DNA products were purified from agarose gels and eluted using the elution kit (Qiagen, 28706). Quick Ligase (NEB, M2200L) was used in all ligation reactions. PCR reactions were carried using Q5 Hot Start Polymerase (NEB, M0493S).

### Cloning IRES under CMV promoter control

Inactivated BVDV-NADL virus was obtained from the laboratory of Paul Walz (Veterinary School, Auburn University). Viral RNA was extracted from 10^6^ viral particles following Ribozol extraction protocol (VWR Life Science, 97064-950). 250 µl of virus suspension was mixed with 3 volumes of Ribozol and incubated for 5 min at room temperature. With an addition of 200 µl chloroform, the sample was shaken for 15 seconds (sec) and incubated for 15 minutes (min) at room temperature. Phases were separated by centrifugation at 12,000 × g for 15 min in a cold microfuge. 600 µl aqueous phase was purified using Zymo Clean & Concentrator-25 (Zymo, R1018). Purified RNA (∼1.2 µg) was used as a template for the synthesis of 461 bp DNA fragment coding for the 5’ UTR sequence and 25 codons of the first viral gene Npro. The oligonucleotide primers were designed using BVDV-NADL. cDNA was synthesized using 40 ng of viral DNA and 1 µM reverse primer IRES-NADL-rev in 20 µl reaction with SuperScript IV reverse transcriptase (ThermoFisher, 18090010). 461 bp DNA fragment was amplified using IRES-NADL forward and reverse primers and Q5 Hot Start polymerase (NEB, M0493S). The second round of PCR amplification extended the 461 bp fragment to include the KpnI restriction site at the 5’ end and the BamHI site at the 3’ end. Amplification with primer pair T7-IRES-for-3 and T7-NADL-rev yielded 509 bp DNA fragment, which was then digested with KpnI-HF (NEB, R3142S) and BamHI-HF (NEB, R3136S). The final 463 bp fragment was ligated with the pcDNA3.1(+)circ RNA Mini Vector (Addgene, 60648) digested with KpnI-HF and BamHI-HF. Clones were selected and maintained in E.coli strain NEB5-alpha (NEB, C2987I). All constructs were verified by Sanger sequencing (Eurofins). New plasmid was designated pIW-IRES.

### Cloning gene for EGFP protein downstream from IRES

Plasmid pIRES-EGFP-puro (Addgene, 45567-DNA.cg) was digested with BstXI (NEB, R0113S) and BsrGI (NEB, R0575S), and 710 bp DNA fragment coding for EGFP protein was prepared for ligation. Plasmid pIW-IRES was digested with BamHI-HF (NEB, R3136S) and ApaI (NEB, R0114S). Insertion of the BstXI-BsrGI fragment with EGFP between BamHI and ApaI sites on the core plasmid was performed using the following adapters: A-5’EGFP-adapt-top and B-5’EGFP-adapt-bott for the 5’ end of the insert plus C-3’EGFP-top and D-3’EGFP-bott for the 3’ end of the insert. The 5’ end adapter pair added an EcoRI restriction site, while the 3’ end pair added an EcoRV site. Double digestion with EcoRI (NEB, R0101S) and EcoRV (NEB, R0195S) was used for the initial screening of clones. All putative correct clones were verified by sequencing. New plasmid was designated pIW-IRES-EGFP.

### Insertion of eMS2 hp at 3’end of IRES

The design of extended hairpin binding bacteriophage MS2 protein was described previously. Plasmid pIW-IRES-EGFP was linearized with BamHI-HF (the 3’ end of IRES) and BstXI (the 5’ end of EGFP). Gblock spanning the 3’ end of IRES, extended protein MS2-binding hairpin (eMS2hp), and the 5’ portion of EGFP were purchased (IDT) and amplified with primers R17L-for and R17L-rev gblock was digested with BamHI and BstXI and ligated to linearized pIW-IRES-EGFP. The sequence of plasmid pIW-IRES(eMS2hp)-EGFP was verified by sequencing.

### RNA isolation and purification

RNA was extracted with TRIzol reagent (ThermoFisher Scientific, 15596018), followed by the RNA Clean and Concentrator kit (Zymo Research, 76020-604) and DNase treatment provided with the kit according to manufacturer’s protocols. The RNA was eluted in 15 µl RNAse free water and stored at − 80 °C.

### *In vitro* transcription

Plasmid pIW-IRES/eMS2hp-EGFP was linearized either with EcoRI for synthesis of IRES/eMS2hp, or with EcoRV for synthesis of IRES/eMS2-EGFP-mRNA. 100 µl transcription reaction consisted of 2.5 µg linearized plasmid, 200 mM HEPES-KOH, 7.5, 30 mM MgCl_2_, 40 mM dithiothreitol (DTT), 1 mM spermidine, 7.5 mM NTP, SUPERNase-In (Ambion) – 20 U/100 µl reaction, inorganic pyrophosphatase (NEB, M0361S) - 0.1 U/100 µl reaction and 100 µg/ml T7 RNA polymerase. The reactions were incubated for 2 hours at 37 °C with gentle rotation.

### BVDV IRES RNA SHAPE-MaP

The sequence corresponding to the BVDV IRES was divided into two overlapping zones (nts 1-283 and 217-504) that were reverse transcribed and amplified with zone-specific RT and PCR primers (Supplementary Table S2). 5 pmol of BVDV IRES *in vitro* transcript (either short, 560 nts or long, 1296 nts) was heated at 95 °C for 3 min and slowly cooled to 4 °C (0.1 °C/sec). The *in vitro* transcripts were folded in a final volume of 36 µl in 1X folding buffer containing 50 mM Tris–HCl pH 8.0, 100 mM NaCl, 5 mM MgCl_2_, and incubated at 37 °C for 15 min. The folded transcripts were divided into two reactions: positive and negative (18 µl each) with one sample treated with 2 µl of 100 mM 1-methyl-7-nitroisatoic anhydride (1M7) (DC Chemicals, 73043-80-8) for positive (+) reaction (final 1M7 concentration of 10 mM), and the second sample treated with 2 µl dimethyl sulfoxide (DMSO) for negative (-) reaction. Both tubes were incubated at 37 °C for 5 min to facilitate RNA modification. Additionally, the denaturing control (DC) was prepared and included 5 pmol of transcript in 3 µl TE-like buffer (0.1 mM EDTA, 10 mM Tris-HCl, pH 7.5), 5 µl formamide and 1 µl denaturing buffer (1 M HEPES, pH 8, 0.5M EDTA), that were mixed and incubated at 95 °C. To 9 µl of the denatured transcript, 1 µl of 100 mM 1M7 was added and incubated at 95 °C for 1 min. All samples were purified using ethanol precipitation with 5 mg/ml glycogen (ThermoFisher, AM9510). After modification, RNA was reverse transcribed at 42 °C for 3 hours using zone-specific RT primers in the SHAPE-MaP buffer, including 50 mM Tris pH 8, 75 mM KCL, 10 mM dithiothreitol (DTT), 6 mM MnCl_2_ (Sigma-Aldrich, 7773-01-5), 0.7 mM dNTPs with 10 U SuperScript II reverse transcriptase (Invitrogen, 18064022). The obtained cDNA products were purified using Mag-Bind Total Pure NGS beads (Omega Bio-Tek, M1378-01). Each sample was amplified separately by two PCR reactions using Q5 high-fidelity polymerase (NEB, M0149S) and zone-specific PCR primers. PCR 1 included 15 cycles: 98 °C, 30 sec; 98 °C, 10 sec; 55 °C, 30 sec; 72 °C, 30 sec; 72 °C, 2 min. The obtained products were checked on 0.8% agarose gel, purified using Mag-Bind Total Pure NGS beads, and subjected to the second round of PCR amplification using 15 cycles: 98 °C, 30 seconds; 98 °C, 10 sec; 55 °C, 30 sec; 72 °C, 30 sec; 72 °C, 2 min. The final libraries were purified on 2% preparative agarose gel by Monarch DNA Gel Extraction kit (NEB, T1020L). The concentration of each library was measured by performing quantitative PCR (qPCR) using PerfeCTa® NGS Quantification Kit (Quanta Biosciences 10029-558) and following the manufacturer’s protocol. Purified amplicon libraries were pooled and sequenced on Illumina MiSeq Instrument.

### SHAPE-MaP data deconvolution

The dataset obtained from Illumina MiSeq was saved into separate FASTQ files for each of the respective zones and reactions. A custom Python script ‘trimmer.py’ was used to trim raw sequences and remove sequencing artifacts. The FASTQ files corresponding to the (+), (-), and (DC) reactions were concatenated and processed by ShapeMapper v1.2 (Busan and Weeks 2018) to obtain mutation counts, sequence read depth, and normalized reactivity values in graphical and CSV format, which help detect potential experimental issues. SuperFold v1.1 (Smola et al. 2015b) was utilized to determine the partition function, minimum free energy, and structural analysis. Shannon entropy was computed from base-pairing probabilities analyzed during SuperFold partition function calculation with a partition Window size of 1200 nts and step size of 100 nts. The local median SHAPE reactivity and Shannon Entropy were calculated over a 55 nt sliding window with a fold step size of 300. RNAstructure v6.0 (Bellaousov et al. 2013) was used for generating the secondary structure models by incorporating SHAPE reactivities as pseudo-energy constraints using slope and intercept values of 2.6 and -0.8, respectively. To determine the presence of a pseudoknot in the IRES RNA, we incorporated the reactivity data into the SHAPEknot v5.8.1 algorithm (34) with the default parameters of SHAPEintercept = -0.8 and SHAPEslope = 2.6. VARNA applet v3-93, ViennaRNA’s RNAplot v2.4.17 and R-chie webserver (Tsybulskyi et al. 2012) were used to visualize RNA secondary structures and arc plots.

### Comparative genomics screen within the genus Pestivirus

Complete genomes of all ICTV-listed Pestivirus species (A-K), as well as two unclassified Pestiviruses, i.e., Linda virus and Norway rat pestivirus were downloaded from the NCBI Genbank database. The cmsearch tool from the Infernal suite v1.1.4 (Nawrocki and Eddy 2013) was used to screen the Rfam covariance model RF00209 (Pestivirus internal ribosome entry site) against the library of 13 Pestivirus genomes, yielding structurally homologous elements within the 5’ UTR in each genome (Supplementary Table S3). These were aligned with mafft v7.475 (Katoh and Standley 2013) and subjected to consensus structure prediction with RNAalifold v2.4.17 (Bernhart et al. 2008) from the ViennaRNA Package v2.4.17 (Lorenz et al. 2011). As RNAalifold cannot directly predict structures that contain pseudoknots, a consensus structure was predicted in two steps. First, a nested, i.e., pseudoknot-free consensus structure was predicted, thereby constraining the regions involved in pseudoknot formation to being unpaired. In a second step, only the pseudoknot was allowed to form, with all other base pairs constrained to being unpaired. A combination of the two predictions yielded the anticipated consensus structure that involves an H-type pseudoknot. Statistical significance of covariation was assessed with R-scape v1.5.10 (Rivas et al. 2016).

### Three-dimensional (3D) structure predictions of BVDV IRES RNA

RNA tertiary structure predictions were performed on a trimmed 106 nt construct that encompasses the portion of the BVDV1-NADL IRES that is involved in pseudoknot formation. The construct was verified to form the expected H-type pseudoknot with pKiss v2.2.13 (Janssen and Giegerich 2015). To test whether the predicted H-type pseudoknot is sterically feasible in 3D, we used ernwin v1.0.1, a tool for sampling coarse-grained 3D structures from fragments of known RNA tertiary structures. Moreover, we performed 3D structure predictions with SimRNA v3.20. This Monte Carlo sampling approach uses a five-bead system per nucleotide to model a coarse-grained representation of RNA structure and employs an empirically derived knowledge-based potential. SimRNA was run for 16 million iterations in replica-exchange mode. The resulting trajectories were subjected to a clustering procedure, thereby filtering sets of similar structures in terms of root mean square deviation (RMSD) within one percent of all trajectories with the lowest energy. Two sets of low-energy candidate structures, comprising 17 and 21 structures, were then evaluated for specific helix angles with forgi v2.0 (Thiel et al. 2019), a library for analyzing RNA tertiary structures. The coarse-grained representations from ernwin and SimRNA, together with the structure probing data, allowed us to assess the feasibility and the structural characteristics of the detected pseudoknot. RNA 3D structures were visualized in PyMol v2.4.2 (Yuan et al. 2017).

## DATA AVAILABILITY

Custom R and Python scripts for data analysis are available at https://github.com/jzs0165

## ACCESSION NUMBERS

The high-throughput sequencing data generated in this study are deposited into SRA databases with bio-sample accession number: PRJNA725417, BVDV-NADL accession number: NC_011461.

## SUPPLEMENTARY DATA

Supplementary Data are available for this article.

## ACKNOWLEDGEMENT

We would like to thank the Alabama Agricultural Experiment Station, Hatch Funding Program for the funding. J.S.S and D.G. are also supported by start-up funds from the Department of Biological Sciences, College of Science and Mathematics, and Office of the Vice President for Research, Auburn University. This work was partly funded by the Austrian science fund (FWF) Doctoral College W 1207 “RNA Biology”.

## Author contributions

“Conceptualization, J.S.S., J.W., M.T.W.; methodology, D.G., I.W., M.T.W., I.K.B, I.L.H.; formal analysis, J.S.S., D.G., J.W., M.T.W., I.K.B, I.L.H, D.G.; writing— original draft preparation, J.S.S., D.G., M.T.W.; writing—review and editing, J.S.S, M.T.W., J.W.; visualization, D.G, I.K.B., M.T.W.; supervision, J.S.S.; project administration, J.S.S, J.W..; funding acquisition, J.S.S., J.W., I.L.H. All authors have read and agreed to the published version of the manuscript.”

